# LTR_FINDER_parallel: parallelization of LTR_FINDER enabling rapid identification of long terminal repeat retrotransposons

**DOI:** 10.1101/722736

**Authors:** Shujun Ou, Ning Jiang

## Abstract

**Summary:** Annotation of plant genomes is still a challenging task due to the abundance of repetitive sequences, especially long terminal repeat (LTR) retrotransposons. LTR_FINDER is a widely used program for identification of LTR retrotransposons but its application on large genomes is hindered by its single threaded processes. Here we report an accessory program that allows parallel operation of LTR_FINDER, resulting up to 8,500X faster identification of LTR elements. It takes only 72 minutes to process the 14.5 Gb bread wheat (*Triticum aestivum*) genome in comparison to 1.16 years required by the original sequential version.

**Availability:** LTR_FINDER_parallel is freely available at https://github.com/oushujun/LTR_FINDER_parallel.

## 1. Introduction

Transposable elements (TEs) are the most prevalent components in eukaryotic genomes. Among different TE classes, long terminal repeat (LTR) retrotransposons, including endogenous retroviruses (ERVs), is one of the most repetitive TEs due to their high copy numbers and large element sizes (Ou and Jiang, 2018). LTR retrotransposons are found in almost all eukaryotes including plants, fungi, and animals, but are most abundant in plant genomes (Bennetzen and Wang, 2014). For example, LTR retrotransposons contribute more than 65% and 70% to the genomes of bread wheat (*Triticum aestivum*) and maize (*Zea mays*), respectively (Ou and Jiang, 2018).

Annotation of LTR retrotransposons relies primarily on *de novo* approaches due to their highly diverse terminal repeats. For this purpose, many computational programs have been developed in the past two decades. LTR_FINDER is one of the most popular LTR search engines and the prediction quality out-performs counterpart programs (Ou and Jiang, 2018). However, LTR_FINDER runs on a single thread and is prohibitively slow for large genomes with long contigs, preventing its application in those species. In this study, we applied the “divide and conquer” approach to simplify and parallel the annotation task for the original LTR_FINDER and observed an up to 8,500 times speedup for analysis of known genomes.

## 2. Methods

We hypothesized that complete sequences of highly complex genomes may contain a large number of complicated nested structures that exponentially increase the search space. To break down these complicated sequence structures, we split chromosomal sequences into relatively short segments (1 Mb) and executes LTR_FINDER in parallel. We expect the time complexity of LTR_FINDER_parallel is *O*(n). For highly complicated regions (i.e., centromeres), one segment could take a rather long time (i.e., hours). To avoid extended operation time in such regions, we used a timeout scheme (300 seconds) to control for the longest time a child process can run. If timeout, the 1 Mb segment is further split into 50 Kb segments to salvage LTR candidates. After processing all segments, the regional coordinates of LTR candidates is converted back to the genome-level coordinates for the convenience of downstream analyses.

LTR_FINDER_parallel is a Perl program that is ready on the go and does not require any form of installation. We used the original LTR_FINDER as the search engine which is binary and also installation free. Based on our previous study (Ou and Jiang, 2018), we applied the optimized parameter for LTR_FINDER (-w 2 -C -D 15000 -d 1000 -L 7000 -l 100 -p 20 -M 0.85), which identifies long terminal repeats ranging from 100 - 7,000 bp with identity ≥ 85% and interval regions from 1 - 15 Kb. The output of LTR_FINDER_parallel is convertible to the popular LTRharvest (Ellinghaus, et al., 2008) format, which is compatible to the high-accuracy post-processing filter LTR_retriever (Ou and Jiang, 2018).

## 3. Results

To benchmark the performance of LTR_FINDER_parallel, we selected four plant genomes with sizes varying from 120 Mb to 14.5 Gb, which are *Arabidopsis thaliana* (version TAIR10) (Arabidopsis Genome Initiative, 2000), *Oryza sativa* (rice, version MSU7) (International Rice Genome Sequencing, 2005; Kawahara, et al., 2013), *Zea mays* (maize, version AGPv4) (Jiao, et al., 2017), and *Triticum aestivum* (wheat, version CS1.0) (International Wheat Genome Sequencing, et al., 2018), respectively. Each of the genomes was analyzed both sequentially (1 thread) and in parallel (36 threads) with wall clock time and maximum memory recorded.

Using our method, we observe 5X - 8,500X increase in speed for plant genomes with varying sizes (Table 1). For the 14.5 Gb bread wheat genome, the original LTR_FINDER took 10,169 hours, or 1.16 years, to complete, while the multithreading version completed in 72 minutes on a modern server with 36 threads, demonstrating an 8,500X increase in speed (Table 1). Even we analyzed each wheat chromosome separately, the original LTR_FINDER still take 20 days in average to complete. Among the genomes we tested, the parallel version of LTR_FINDER produced slightly different numbers of LTR candidates when compared to those generated using the original version (0% - 2.73%; Table 1), which is likely due to the use of the dynamic task control approach for processing of heavily nested regions. Given the substantial speed improvement (Table 1), we consider the parallel version to be a promising solution for large genomes.

**Table 1.**
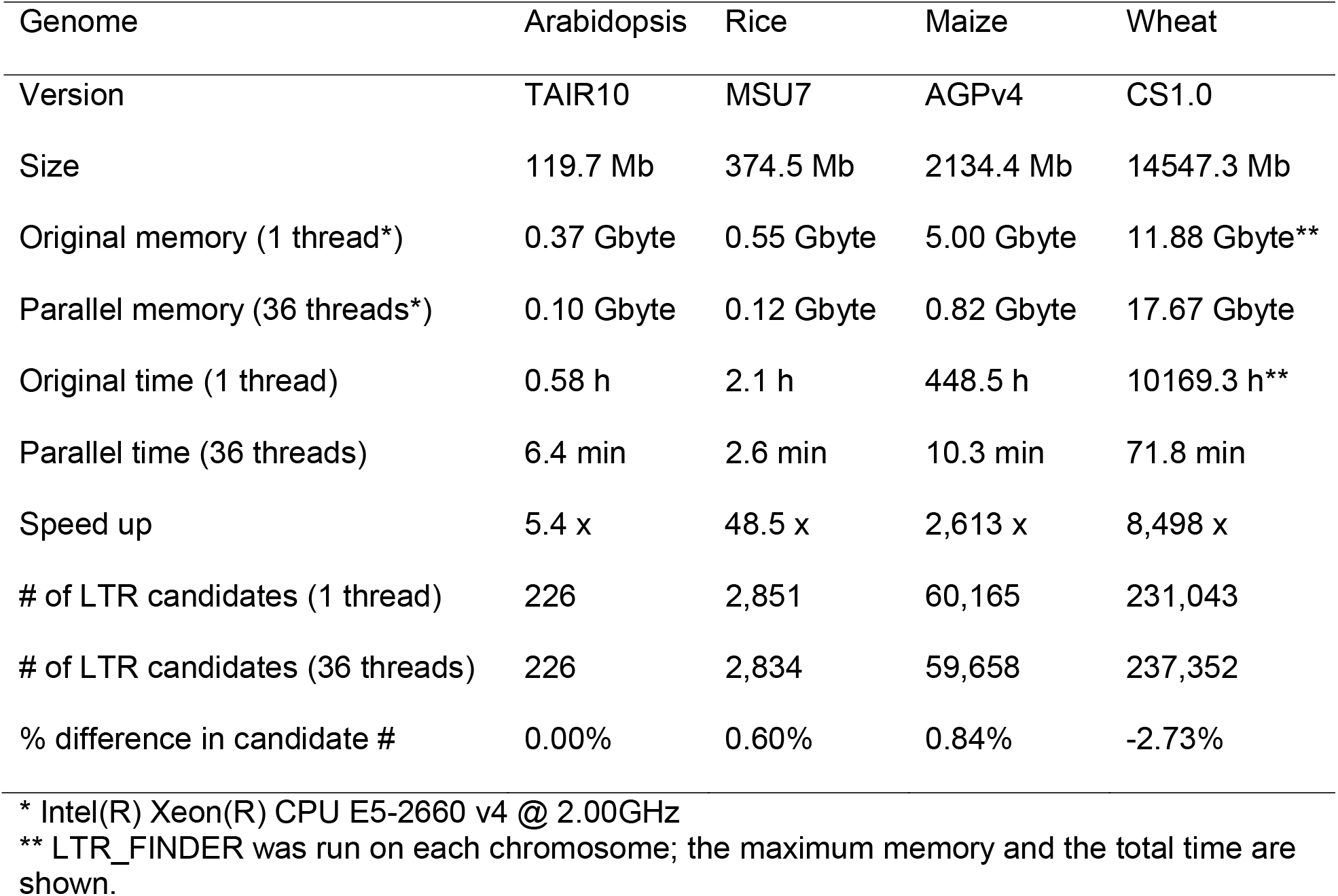
Benchmarking the performance of LTR_FINDER_parallel.

| Genome | Arabidopsis | Rice | Maize | Wheat |
| --- | --- | --- | --- | --- |
| Version | TAIR10 | MSU7 | AGPv4 | CS1.0 |
| Size | 119.7 Mb | 374.5 Mb | 2134.4 Mb | 14547.3 Mb |
| Original memory (1 thread*) | 0.37 Gbyte | 0.55 Gbyte | 5.00 Gbyte | 11.88 Gbyte** |
| Parallel memory (36 threads*) | 0.10 Gbyte | 0.12 Gbyte | 0.82 Gbyte | 17.67 Gbyte |
| Original time (1 thread) | 0.58 h | 2.1 h | 448.5 h | 10169.3 h** |
| Parallel time (36 threads) | 6.4 min | 2.6 min | 10.3 min | 71.8 min |
| Speed up | 5.4 x | 48.5 x | 2,613 x | 8,498 x |
| # of LTR candidates (1 thread) | 226 | 2,851 | 60,165 | 231,043 |
| # of LTR candidates (36 threads) | 226 | 2,834 | 59,658 | 237,352 |
| % difference in candidate # | 0.00% | 0.60% | 0.84% | -2.73% |

\* Intel(R) Xeon(R) CPU E5-2660 v4 @ 2.00GHz
\*\* LTR\_FINDER was run on each chromosome; the maximum memory and the total time are shown.

